# Ecosystem gross primary productivity after autumn snowfall and melt events in a mountain meadow

**DOI:** 10.1101/2022.01.25.477606

**Authors:** P. C. Stoy, A. M. Khan, K. Van Dorsten, P. Sauer, T. Weaver, E. N. J. Brookshire

## Abstract

Vegetation productivity is increasing across much of the U.S. Northern Great Plains but is decreasing in some nearby Northern Rocky Mountain grasslands due to increases in aridity. Mountain grasslands support critical ecosystem services that are under threat from ongoing land use and climate changes, and it is important to understand their function across all changing seasons. Observing the full range of montane ecosystem productivity is challenging because site access is often difficult during the “shoulder seasons” in spring and autumn if the snowpack is not fully developed or degrading. It is unclear if decreases to montane grassland productivity from drying autumns can be offset in part by late-season green-ups after precipitation events. These include the snowfall/snowmelt periods that often characterize the summer-to-winter transition in the Northern Rockies. Here, we quantify the ecosystem carbon uptake that occurs after snowfall and melt in climatological autumn (September, October, and November) in a montane grassland in Montana, USA using a combination of eddy covariance, phenological camera, and remote sensing analyses. Carbon dioxide flux follows a diurnal pattern after autumn snowmelt events despite overall ecosystem losses of C, suggesting that post-snowmelt photosynthesis helps dampen C loss during autumn and provides fresh photosynthate to support ecosystem functioning. Light-saturated photosynthesis after two snow events was not different than before snowfall (∼6 µmol CO_2_ m^-2^ s^-1^ in 2016 and ∼2.5 µmol CO_2_ m^-2^ s^-1^ in 2017); observations are consistent with the notion that canopy photosynthesis is resistant, rather than resilient, to the first snow disturbances. MODIS observations also suggest that post-snowfall increases in NDVI can occur but do not happen every year, such that late-season photosynthesis is not a reliable source of fresh photosynthate. These late-season carbon uptake events likely play a small role in the annual ecosystem carbon balance but may be disproportionately important for organisms faced with dwindling late-season forage and regrowth in spring. Future efforts should seek to understand the community and ecosystem consequences of vegetation functioning during autumn as part of an expanded effort to understand phenological changes during this under-studied and changing time of year.

## Introduction

North American grassland productivity is predicted to increase due to vegetation responses to future climatic variability (Hufkens et al. 2016) despite well-documented increases in aridity (Novick et al. 2016). Widespread greening has already been observed across the Northern Great Plains (Brookshire et al. 2020), but the growing season across many surrounding montane regions has become shorter due to hydrologic stress (Wood et al. 2021), with implications for biogeochemical cycling and ecosystem functioning. Late-season productivity, for example, has declined by over 50% since the late 1960s at a well-studied montane grassland site in the US Northern Rocky Mountains due to increases in aridity (Brookshire and Weaver 2015), despite regional climate changes that are often favorable for plant growth (Bromley et al. 2020). These observations suggest that Northern Rocky Mountain grasslands are responding differently than Northern Great Plains grasslands to recent climate changes and may continue to do so.

Montane grasslands support diverse biological communities (Körner and Spehn 2019) and provide critical ecosystem services (Körner 2004) that are under threat from changes to land use and climate (Egan and Price 2017) including snow dynamics (Harpold et al. 2012). Carbon sequestration is a critical ecosystem service that may be substantial in montane grasslands (Reed et al. 2021) but is highly variable across space and time – due in part to climate and management pressures (Wohlfahrt, et al. 2008; Rogiers et al. 2004; Schmitt et al. 2010; Tenhunen et al. 2009) – which may result in net annual CO_2_ losses (Rogiers et al. 2008). Regardless of net annual carbon efflux, the documented pronounced late season declines of vegetation productivity in montane grasslands of the northern Rocky Mountains is cause for concern for the organisms that rely on fresh vegetation for overwintering success (Hurley et al. 2014, 2017; Loe et al. 2020).

Despite large declines in biomass during the late growing season before snowfall in a mountain meadow in Montana (Brookshire and Weaver 2015), research at the site and communication with land managers has led to the speculation that precipitation pulses – often from snowfall and subsequent ablation (hereafter “melt”) – support a “recovery” of production during the transition from the dry summer season to winter. This notion is consistent with a hypothesis that montane grassland photosynthesis is *resilient* to late-season snow events and can quickly re-establish previous function after melt, much in the same way that grass photosynthesis tends to re-establish quickly following harvesting (Novick et al. 2004; Stoy et al. 2008), grazing (Parsons et al. 1983), and heat waves (Hooveret al. 2014). Alternately, canopy photosynthesis in mountain meadows could simply be *resistant* to snow disturbances following a growing body of cold-season research that demonstrates that photosynthesis can occur under snow (Starr and Oberbauer 2003; Street et al. 2012; Lundell et al. 2008) and the notion that grass productivity is resistant to minor disturbances, including for example mild drought (Batbaatar et al. 2021). In this scenario, ecosystem carbon uptake would not meaningfully differ before and after snow events and ecosystem carbon uptake would have little response to snow apart from the limitations to photosynthesis caused by cold conditions and diminished light availability under the snowpack. Most research to date has characterized the rapid re-establishment of photosynthesis after snowmelt in spring (Julitta et al. 2014; Vitasse et al. 2017; Wohlfahrt et al. 2008), with fewer studies of ecosystem carbon uptake in response to short-duration snow events during the autumn-to-winter transition.

Ecological research in montane ecosystems during autumn and the transition from the vegetative growing season to winter is lacking, in part due to the difficult travel conditions created by snowfall and the subsequent melt periods that make roads impassable and over-snow transport onerous. This has hindered our understanding of ecosystem functioning during this critical and changing period (Piao et al. 2008), especially as compared to its spring counterpart (Gallinat et al. 2015). To help remedy this situation for the case of montane grasslands, we studied the effects of autumn snowfall and melt periods on ecosystem carbon uptake and loss in a montane grassland in Montana, USA that has been studied for decades. Specifically, we characterize the surface-atmosphere exchange of carbon dioxide and energy, greenness, and the normalized difference vegetation index (NDVI) before, during, and after winter snow accumulation and melt events in September, October, and November – climatological autumn – using eddy covariance, phenocam, and remote sensing measurements and ask if observations follow a pattern of ecological resilience or resistance to autumnal snow.

## Methods

### Site information

Observations were made at the Bangtail Study Area (Brookshire and Weaver 2015; Weaver and Collins 1977), which hosts the Bangtail Mountain Meadow eddy covariance research site (US-BMM). It is located 24 km northeast of Bozeman, MT, USA on a windswept ridge at 2324 m above sea level at 45.7830 N, 110.7776 W. The site served as the high elevation grassland biome site for the International Biological Program (1964-1974) and hosts the longest-active snow manipulation experiment of which we are aware (Brookshire and Weaver 2015; Yano et al. 2015). The mean annual temperature is 2.3 ℃ and the mean annual precipitation is 981 mm as estimated by *daymet* (Thornton et al. 2014). Vegetation is dominated by Idaho fescue (*Festuca idahoensis*) with increasing cover of the invasive Kentucky bluegrass (*Poa pratensis*) and soils are loamy-typic cryoborolls (Yano et al. 2015; Weaver and Collins 1977). The study area has been fenced against cattle since the 1930s but is available for forage by wild large ungulates (e.g. mule deer (*Odocoileus hemionus*) and elk (*Cervus canadensis*)) and smaller subalpine animals including marmot (*Marmota caligata*), deer mouse (*Peromyscus maniculatus*), northern pocket gopher (*Thomomys talpoides*), and more.

### Eddy covariance and micrometeorological instrumentation

A 3-m tall tripod tower was installed on September 30, 2016, to host eddy covariance and micrometeorological instrumentation (Figure A1). Measurements became available on October 1, 2016. Eddy fluxes were measured using a three-dimensional sonic anemometer (CSAT-3, Campbell Scientific, Logan, UT, USA) mounted at 153 cm above ground level and an enclosed infrared gas analyzer (EC-155 Campbell Scientific) – both recording at 10 Hz to a CR6 datalogger (Campbell Scientific) – as part of a CPEC300 eddy covariance system. Air temperature (TA) and relative humidity were measured using an HMP45C (Vaisala, Vantaa, Finland) at 200 cm and used to calculate the atmospheric vapor pressure deficit (VPD). Incident and outgoing short- and longwave radiation were measured using an NR01 net radiometer (Hukseflux, Delft, The Netherlands) at 176 cm. Outgoing longwave radiation observations were used to estimate the radiometric surface temperature (Tsurf) using the Stefan-Boltzmann equation, assuming a view factor of unity and an emissivity of 0.98 to account for all surfaces observed by the radiometer including grass, bare ground, and snow (Snyder et al. 1998). The shortwave albedo (ALB) was calculated as the ratio of outgoing to incident shortwave radiation, and we use the median of measurements taken between 1100 and 1300 local time to minimize solar zenith angle effects. The normalized difference vegetation index (NDVI) and photochemical reflectance index (PRI) were measured at 176 cm using Spectral Reflectance Sensors (Decagon (now METER), Pullman, WA, USA). We likewise use the median of measurements taken between 1100 and 1300 to calculate daily values. Soil heat flux (G) was measured using two HFP01SC heat flux plates (Hukseflux) at 10 cm below the soil surface. Three CS650 soil temperature (Tsoil) and moisture content (SWC) sensors (Campbell Scientific) were positioned horizontally at 10, 20, and 40 cm below the soil surface to account for a rooting depth observed to be at least 50 cm. An additional CS650 was positioned vertically to make an integrated measurement of the top 30 cm of soil. We present measurements from the sensor at 10 cm to characterize conditions near the surface including potential freezing. An SR50 sonic distance sensor (Campbell Scientific) was pointed downward at 146 cm to measure distance to surface; we use the median observation from each day to characterize vegetation height or snow depth (h). Micrometeorological observations were measured every second, averaged to half-hour periods, and saved to a CR1000 datalogger (Campbell Scientific). A list of abbreviations is presented in Table 1.

**Table 1:** List of abbreviations with units.

| Abbreviation | Definition | Units or value |
| --- | --- | --- |
| ALB | Shortwave albedo | unitless |
| DN | Digital number | unitless |
| $E_0$ | Related to activation energy | $K^{-1}$ |
| $F_c$ | Eddy flux of carbon dioxide | $\mu\text{mol m}^{-2} \text{s}^{-1}$ |
| G | Ground heat flux | $\text{W m}^{-2}$ |
| GCC | Green chromatic coordinate | unitless |
| GPP | Gross primary productivity | $\mu\text{mol m}^{-2} \text{s}^{-1}$ |
| h | Height | m |
| H | Sensible heat flux | $\text{W m}^{-2}$ |
| k | Parameter describing sensitivity of $\beta$ to VPD | $\text{hPa}^{-1}$ |
| LE | Latent heat flux | $\text{W m}^{-2}$ |
| NDVI | Normalized difference vegetation index | unitless |
| NDVI | NDVI from MODIS | unitless |
| PPFD | Photosynthetic photon flux density | $\mu\text{mol m}^{-2} \text{s}^{-1}$ |
| PRI | Photochemical reflectance index | unitless |
| RCC | Red chromatic coordinate | unitless |
| $R_g$ | Global shortwave radiation | $\text{W m}^{-2}$ |
| RGB | Red, green, blue | See DN |
| ROI | Region of interest | $\text{m}^2$ |
| SWC | Soil water content | % |
| $T_0$ | Respiration model constant | $-46.02\text{ }^{\circ}\text{C}$ |
| TA | Air temperature | $^{\circ}\text{C}$ |
| $T_{\text{ref}}$ | Reference temperature | 15 °C |
| $T_{\text{soil}}$ | Soil temperature | °C |
| $T_{\text{surf}}$ | Radiometric surface temperature | °C |
| VPD | Vapor pressure deficit | hPa |
| $\text{VPD}_0$ | VPD threshold for $\beta$ limitation | 10 hPa |
| $\alpha$ | Initial slope of the light-response curve | $\mu\text{mol CO}_2 \text{ J}^{-1}$ |
| $\beta$ | Light response curve at saturation | $\mu\text{mol m}^{-2} \text{ s}^{-1}$ |
| $\beta_0$ | Maximum $\beta$ at non-limiting VPD | $\mu\text{mol m}^{-2} \text{ s}^{-1}$ |
| $\gamma$ | Fitted parameter equivalent to RE | $\mu\text{mol m}^{-2} \text{ s}^{-1}$ |
| $\rho_{\text{NIR}}$ | Near-infrared reflectance | unitless |
| $\rho_{\text{Red}}$ | Red reflectance | unitless |

### Eddy covariance data processing

Half-hourly measurements of the eddy flux of carbon dioxide (Fc), latent heat flux (LE), sensible heat flux (H), and momentum flux were calculated using EddyPro (LiCor, Lincoln, Nebraska, USA). To calculate fluxes, we used double rotation (Kaimal and Finnigan 1994) – rather than the planar fit method – due to rapid changes in canopy and snow height, as well as block averaging, covariance maximization for time lag detection, and high- and low-pass filters (Moncrieff et al. 2005; Moncrieff et al. 1997). Spike removal was performed as described by Vickers and Mahrt (1997) and defined as more than 3.5 standard deviations from the mean mixing ratio for water vapor and carbon dioxide and 5 standard deviations for vertical wind velocity. Fluxes that did not pass the Mauder and Foken (2011) quality control tests were excluded from the analysis. We assumed that storage of carbon dioxide below the 153 cm instrument height was negligible such that Fc approximates the net ecosystem exchange of carbon dioxide (NEE). Open access data from US-BMM are published on the Ameriflux website at https://ameriflux.lbl.gov/sites/siteinfo/US-BMM (Stoy, 2021).

NEE was partitioned into its components, gross primary productivity (GPP) and ecosystem respiration (RE), using REddyProc (Wutzler et al. 2018). We explored flux partitioning methods that use nighttime flux measurements (Reichstein et al. 2005) and both daytime nighttime and daytime flux measurements (Lasslop et al. 2010) to model NEE given that results from these two methods can differ and impact the interpretation of GPP (Reichstein et al. 2012; Stoy et al. 2006).

The partitioning method based on nighttime data develops a relationship between RE and air temperature as described in Reichstein et al. (2005):

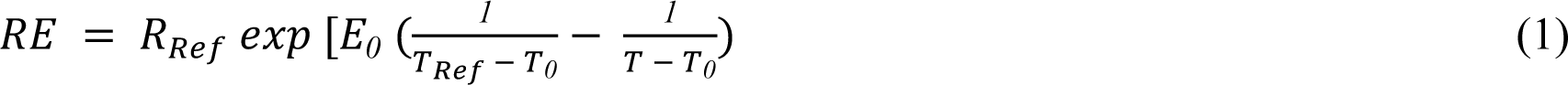

where *R_Ref_* (μmol CO_2_ m^-2^ s^-1^) is the reference respiration at the reference temperature, *T_Ref_* is set to 15 ℃ and *T_0_* is set to −46.02 ℃ (Wutzler et al. 2018; Reichstein et al. 2005; Lloyd and Taylor 1994). Nighttime data are constrained between local sunrise and sunset and when incoming shortwave radiation is less than 10 W m^-2^. *E_0_* is estimated using 15-day windows and aggregated to an annual estimate, and a temporally varying *R_Ref_* is estimated using the annually aggregated *E_0_* (Wutzler et al. 2018). The relationship developed from nighttime data is extrapolated using air temperature measurements during the day to estimate daytime RE and GPP is calculated as: GPP = RE − NEE.

The partitioning method based on nighttime and daytime data estimates NEE using a light response curve following (Lasslop et al. 2010):

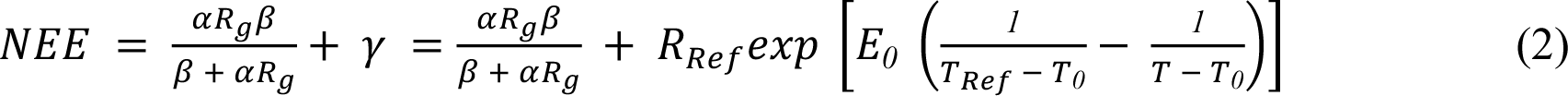

where α (μmol CO_2_ J^-1^) is the canopy light use efficiency before light saturation is reached, β (μmol CO_2_ m^-2^ s^-1^) is the maximum CO_2_ uptake rate at light saturation, *R_g_* is the incoming shortwave radiation at the surface (W m^-2^), and γ (μmol CO_2_ m^-2^ s^-1^) is RE. β in the Lasslop et al. (2010) algorithm is calculated as:

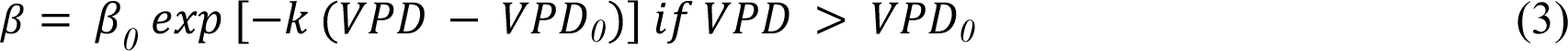

where β_0_ is the maximum CO_2_ uptake rate at light saturation under ideal VPD and β = β;_0_ when VPD ≤ *VPD_0_*. *VPD_0_* is set as 10 hPa. *E_0_* is estimated using nighttime data in moving windows and the resulting estimates are smoothed (Wutzler et al. 2018). Prior estimates for *R_Ref_* are also estimated using nighttime data for the moving window with smoothed *E_0_*. Finally, the parameters (*R_Ref_*, α, β_0_, *k*) are estimated using daytime data for each time window (Wutzler et al. 2018).

The first term on the right-hand side of the equation (2) represents GPP and we fit the equation 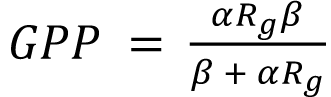 using nonlinear least squares (nls) in R (R Core Team, 2021) to calculate β for periods before and between snow events. We use ANOVA with Tukey’s HSD test to calculate significant differences in β among different measurement periods. Latent heat flux (LE) and sensible heat flux (H) were also gapfilled using REddyProc, which applies mean diurnal sampling to estimate missing values of these quantities (Falge et al. 2001).

### Vegetation indices from Phenocam and MODIS

A 5 MP NetCam SC phenocam (StarDot Technologies, Buena Park, CA) was installed 75 m SW of the eddy covariance tower in August 2015. The phenocam pointed north and captured an area that is typically upwind of the tower given the predominantly westerly winds (Figure 1). Observations were available for November and December 2016 after eddy covariance observations began, and for 2017 until early October.

**Figure 1:**
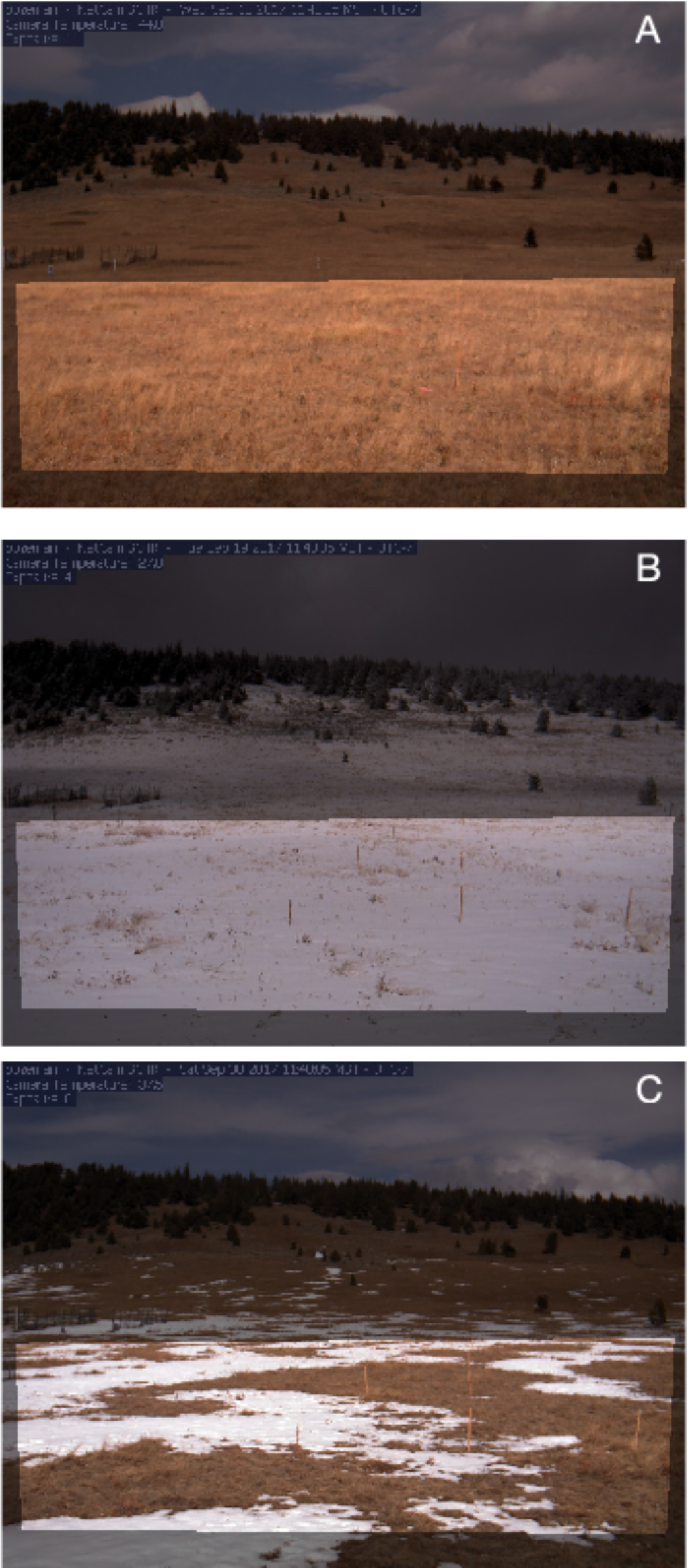
Three representative phenocam images taken near noon local standard time on September 13 (A), September 19 (B), and September 30 (C), 2017. Areas outside the grass region of interest (ROI) are darkened with a translucent mask.

Hourly daytime .jpg images were automatically uploaded to the PhenoCam website (https://phenocam.sr.unh.edu/webcam) where they are freely available. For this analysis we use the phenocam data product of (Seyednasrollah et al. 2019) as described in (Richardson et al. 2018) and (Seyednasrollah et al. 2019). Briefly, the phenocam analysis involves choosing a region of interest (ROI) for the main grass portion of each image as demonstrated for a single image in Figure 1. Digital numbers (DN) for the red, green, and blue (RGB) layers were then extracted from the ROI of each image and used to calculate the green chromatic coordinate (GCC) and red chromatic coordinate (RCC), which have proven effective for describing phenological variability (Richardson et al. 2009, 2007) while minimizing the effects of variable scene illumination, noting that the RCC has proven particularly useful for identifying snow (Liu et al. 2020):

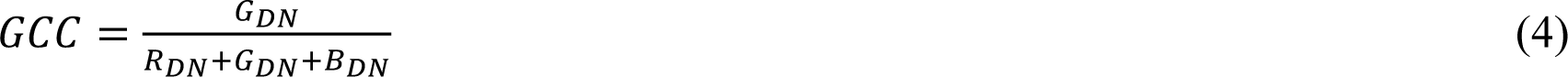

and

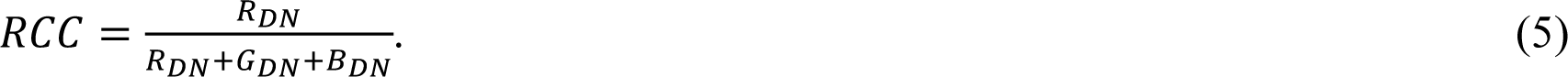

For this analysis we use the daily 90th percentile values of the GCC and RCC as recommended by (Richardson et al. 2018) to minimize the effects of diurnal variability in illumination geometry and its effects on phenocam imagery.

To explore late-season greenness from spaceborne platforms, we calculated NDVI for the pixel corresponding to the tower (45.784884, −110.777995) using the MCD43A4 v006 Terra+Aqua Nadir BRDF-Adjusted Reflectance daily 500 m product (Schaaf and Wang 2015; Campagnolo et al. 2016). The NDVI was calculated as:

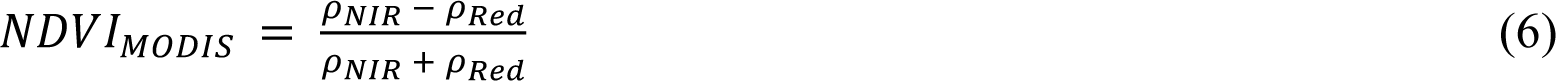

Where ρ_NIR_ is the MCD43A4 surface reflectance in near-infrared wavelengths (841 - 876 nm) and ρ_Red_ is the MCD43A4 surface reflectance in red wavelengths (620 - 670 nm).

## Results

We first present micrometeorological, surface-atmosphere exchange, and radiometric data for the 2016 study period, which we define as October 1 when measurements began until November 12 when flux measurements became unavailable due to power system failure. We follow the 2016 results with the 2017 findings and define a study period of September 10 – four days before first snowfall for consistency with the 2016 study period – until November 12. Measurements were available until December 12, 2017, but with consistent snow cover after October 29 as we detail later. We then discuss measurements made during the summer of 2017 for comparison with autumn measurements and describe MODIS observations for the years prior to, during, and after the eddy covariance study period.

### Observations during autumn, 2016

G decreased from a maximum of 34 W m^-2^ when measurements began on October 1, 2016 (Figure 2) and did not exhibit a diurnal cycle four days later, when Tsurf was equal to or less than 0 ℃, SWC increased from 30% to 35%, ALB increased to values greater than 50%, NDVI decreased to less than 0, and h (here suggesting changes in snow height) increased from 0.12 to 0.20 m (Figure 3). These observations are consistent with snow presence, and we subsequently use micrometeorological, radiometric, and flux observations to determine snow presence or absence (Table 2). We also use phenocam observations to identify snow periods during the parts of 2017 when images were available (e.g., Figure 1).

**Figure 2:**
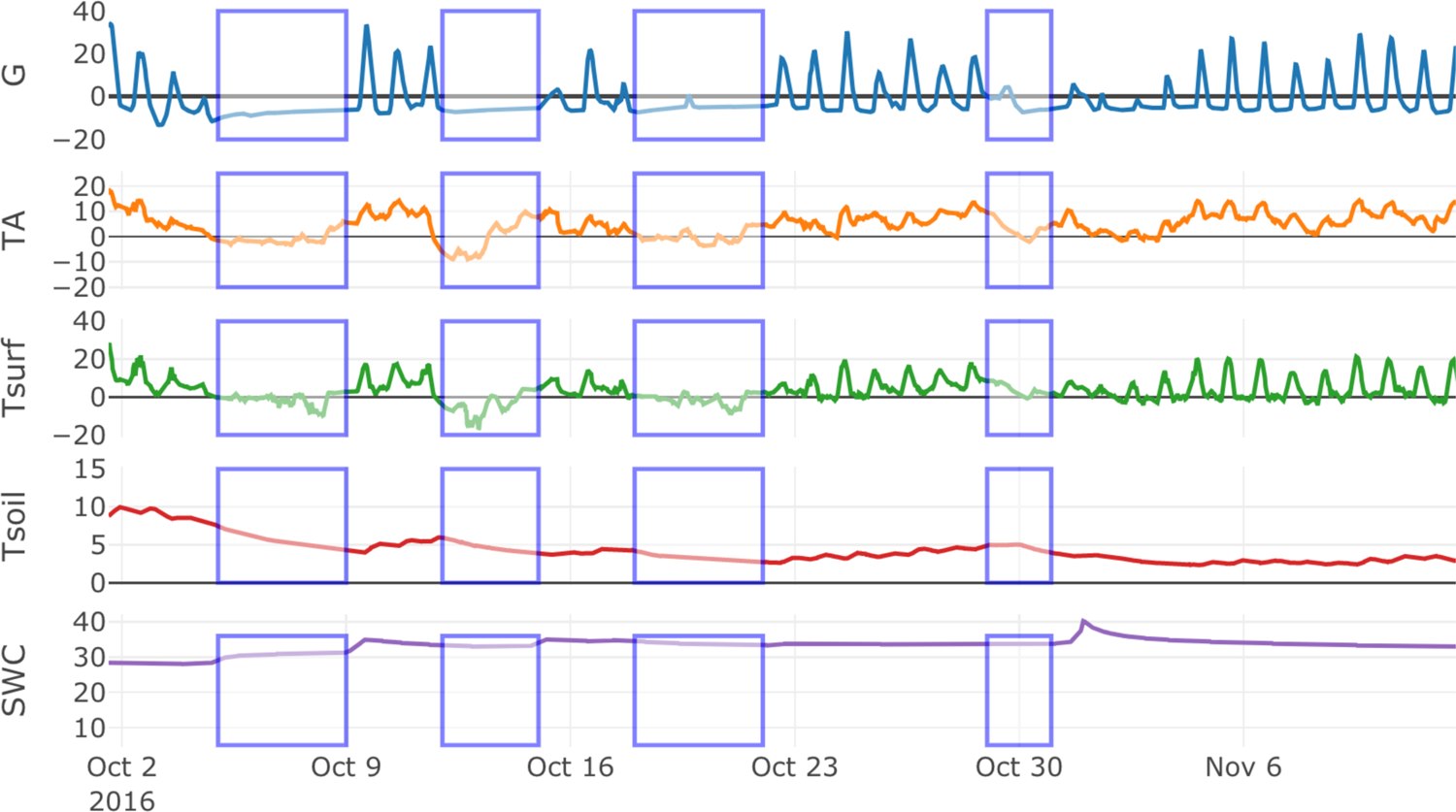
Micrometeorological conditions during the 2016 study period. Periods identified as having snow (Table 2) are denoted by in blue boxes. Units are W m^-2^ for soil heat flux (G) with the convention that positive values indicate heat flux into the soil, ℃ for air temperature (TA), radiometric surface temperature (Tsurf), and soil temperature (Tsoil), and % for volumetric soil water content (SWC).

**Figure 3:**
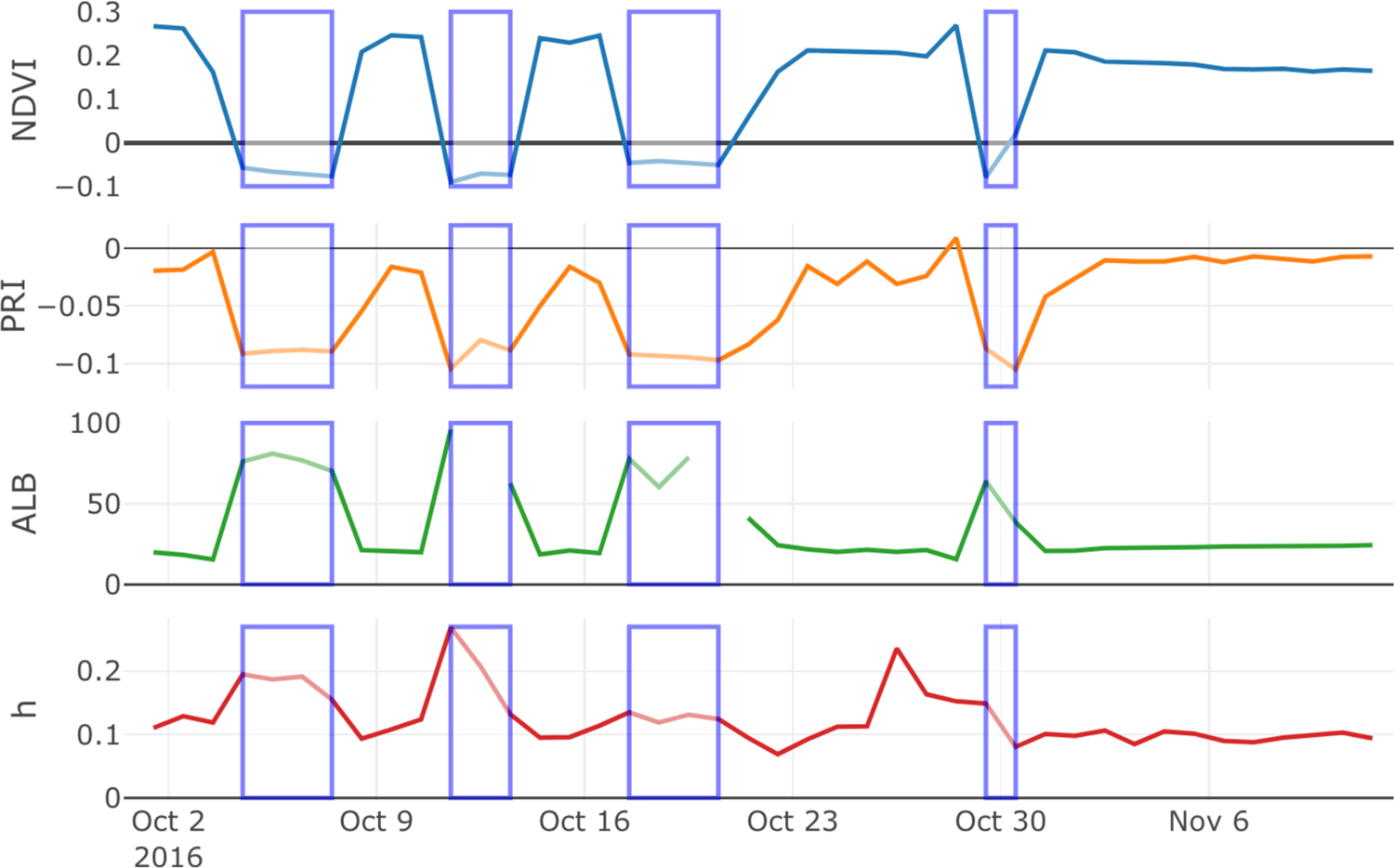
The median normalized difference vegetation index (NDVI), photochemical reflectance index (PRI), and shortwave albedo (ALB) for 11:00 - 13:00 local time during October and early November 2016. Grass or snow height (h, m) was calculated as the median half-hourly sonic distance measurement for each day. Periods identified as having snow (Table 2) are indicated in blue boxes.

**Table 2:** The dates of the before-snow period, snow presence (S) and inter-snow periods (IS), numbered sequentially, at the Bangtail Mountain Meadow (US-BMM) eddy covariance research site in Montana, USA during 2016. Light response curves were not calculated for snow periods despite non-zero GPP (e.g., Figure 5) due to the frequent patchy snow conditions (e.g. Figure 1).

| Event | Dates | $\beta$ ($\mu\text{mol CO}_2 \text{ m}^{-2} \text{ s}^{-1}$) |
| --- | --- | --- |
| Before Snow | Oct. 1<br>Oct. 4 | $6.23 \pm 0.84^a$ |
| S1 | Oct. 5<br>Oct. 8 | - |
| IS1 | Oct. 9<br>Oct. 11 | $7.07 \pm 0.58^a$ |
| S2 | Oct. 12<br>Oct. 14 | - |
| IS2 | Oct. 15<br>Oct. 17 | $6.63 \pm 0.38^a$ |
| S3 | Oct. 18<br>Oct. 21 | - |
| IS3 | Oct. 22<br>Oct. 28 | $4.62 \pm 0.19^b$ |
| S4 | Oct. 29<br>Oct. 30 | - |
| IS4 | Oct. 31<br>Nov. 12* | $3.44 \pm 0.15^c$ |

\* Measurements became unavailable for the rest of the year after Nov. 12, 2016.

NEE exhibited a distinct diurnal signal before the first snowfall event of autumn, 2016, that was dampened when snow was present (Figure 4). The nighttime partitioning approach (Reichstein et al. 2005) estimated that GPP was usually no greater than 2 μmol m^-2^ s^-1^ during peak daytime periods when snow was present. The daytime partitioning approach (Lasslop et al. 2010) frequently estimated a GPP of ∼2 μmol m^-2^ s^-1^ during peak daytime periods when snow was present (Figure A2). From these observations, nighttime partitioning is arguably more conservative for estimating GPP in the presence of snow. We focus the rest of our analysis on GPP and RE values obtained using the Reichstein et al. (2005) nighttime partitioning method.

**Figure 4:**
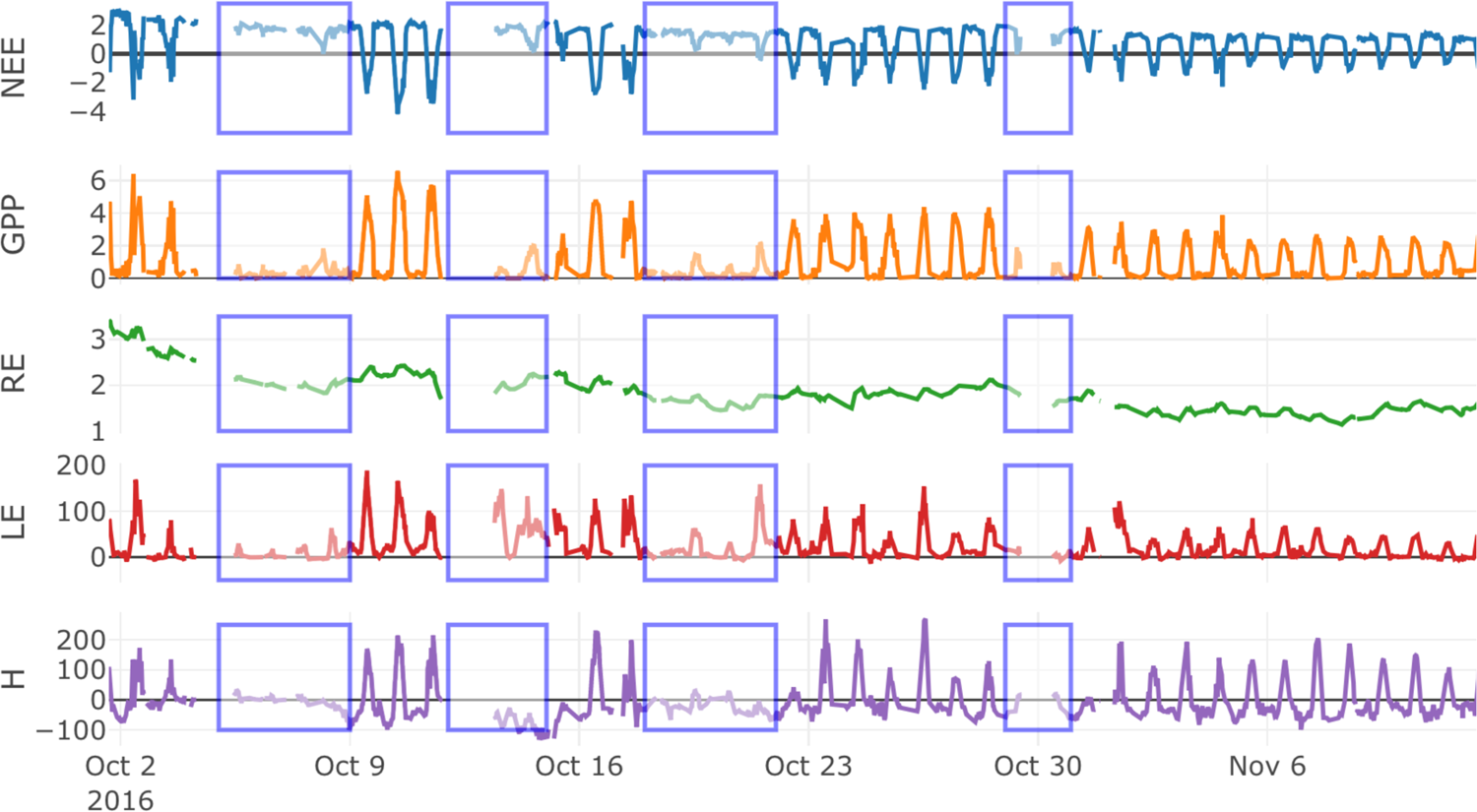
The net ecosystem exchange (NEE), gross primary productivity (GPP), ecosystem respiration (RE), latent heat flux (LE), and sensible heat flux (H) during October and November 2016 at the Bangtail Mountain Meadow, Montana, USA study site. Periods identified as having snow (Table 2) are indicated in blue boxes. Units for NEE, GPP, and RE are (µmol CO_2_ m^-2^ s^-1^) and units for LE and H are W m^-2^.

We identified four snow periods in 2016 (Table 2, Figures 2-4) before measurements became unavailable for the rest of the year on November 12. After the first snowfall, Tsoil was lower and SWC higher than the pre-snow period for the remainder of the 2016 study period. Tsoil remained lower than 6 ℃ after first snowfall for the remainder of the study period, but TA consistently reached 10 ℃ or greater and Tsurf was frequently greater than 20 ℃ during midday periods between snowfall events. Mid-day reflectance measurements (NDVI and PRI) did not differ for the period before snow and the inter-snow periods (Figure 3).

Mid-day GPP decreased from ∼6 μmol m^-2^ s^-1^ before the first snowfall to between 2 and 4 ∼ μmol m^-2^ s^-1^ after the fourth snow event (Figure 4). The light response curve analysis indicated no significant difference in β for the period before snow and first two inter-snow periods, which averaged 6.64 µmol CO_2_ m^-2^ s^-1^ (Figure 5, Table 2). β during the third inter-snow period was significantly lower (4.62 µmol CO_2_ m^-2^ s^-1^) and the fourth significantly lower still (3.44 µmol CO_2_ m^-2^ s^-1^) (Table 2). Respiration decreased from 3 μmol m^-2^ s^-1^ to 1 μmol m^-2^ s^-1^ throughout the 2016 autumn measurement period (Figure 4). As a consequence, the ecosystem was a net sink of CO_2_ (i.e., NEE was negative) during midday of all snow-free periods during the measurement record in 2016 (Figure 4) but was a source of C to the atmosphere during other times of day and lost 50 g C m^-2^ from October 2 until November 12, 2016.

**Figure 5:**
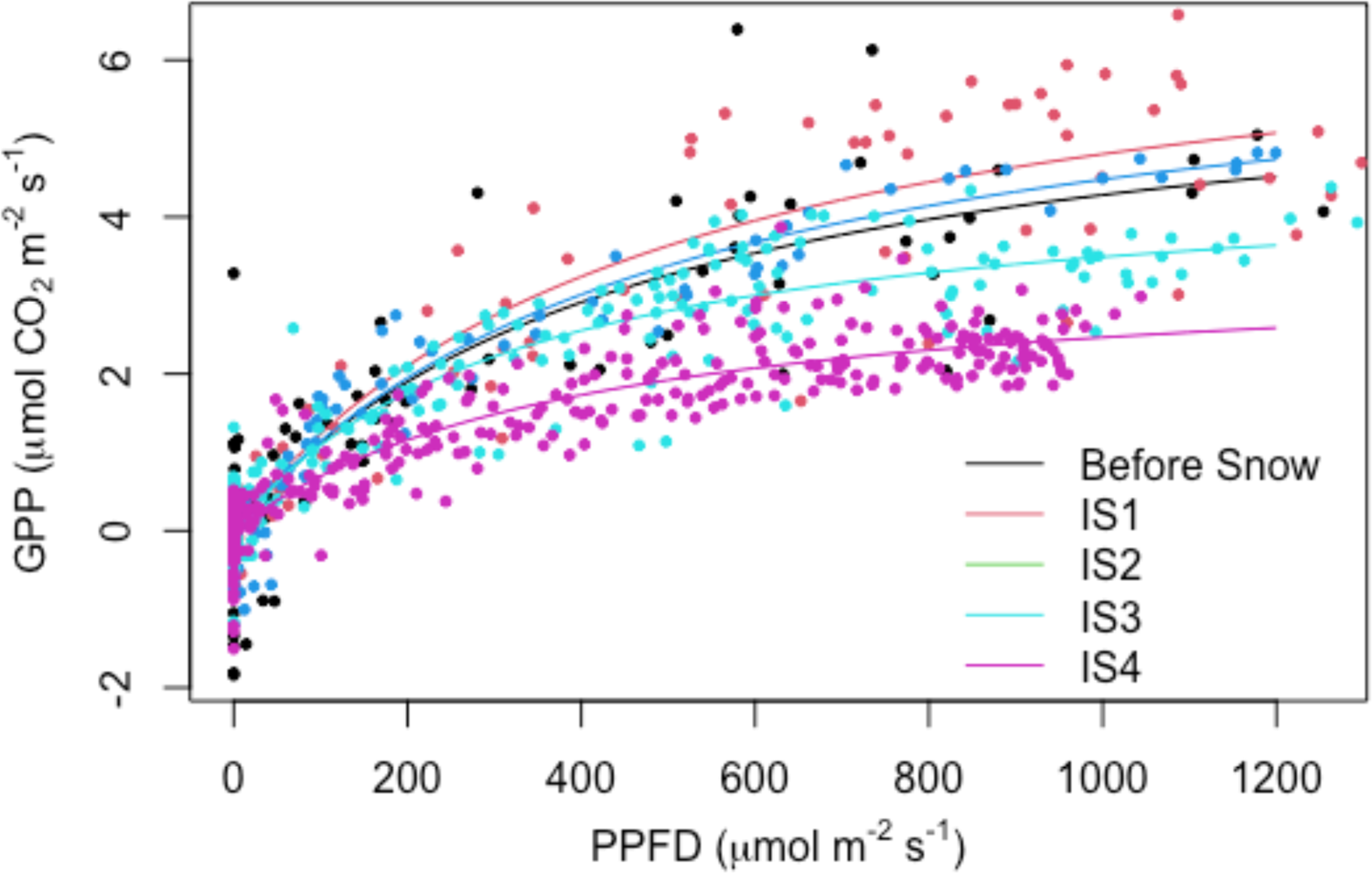
The relationship between GPP and PPFD with fitted light response curves (equation 2) for before snow and inter-snow (IS) periods (Table 1) at the Bangtail Mountain Meadow study site during 2016.

### Observations during autumn, 2017

Autumn, 2017 was characterized by two weeks of snow presence from September 15 until September 29, after which measurements indicated a three-day snow-free period, followed by four days of snow, nearly a week without snow, and additional snow and no-snow periods of less than one week in duration before consistent snow presence began on October 29. We determined these dates using a combination of G, Tsurf, and turbulent flux observations and phenocam observations when available, and we discuss challenges in determining snow presence in the Discussion.

SWC was ∼ 10% before snowfall (Figure 6) – recalling that SWC was ∼30% before snow fell in 2016 (Figure 2) – and increased to 35% after the first 2017 snow event. TA intermittently reached nearly 20 ℃ during the periods between snow in 2017, as did Tsurf, and Tsoil remained above 0 ℃ until November 12 when the study period ended for consistency with data availability during 2016 (Figure 6). As opposed to 2016, NDVI stayed consistently at or below 0 and ALB remained above 50% during the periods between snow presence (Figure 7), which we identified largely using G, Tsurf, and turbulent flux observations as noted. The phenocam was active during the first and second snowfall and melt periods in 2017, and RCC declined sharply from nearly 0.6 to less than 0.35 after the first snow event (Figure 8). RCC recovered to values approaching 0.4 during the first inter-snow period and nearly 0.45 after the second snow period after which measurements became unavailable. GCC was less sensitive to snow presence and absence (Figure 8).

**Figure 6:**
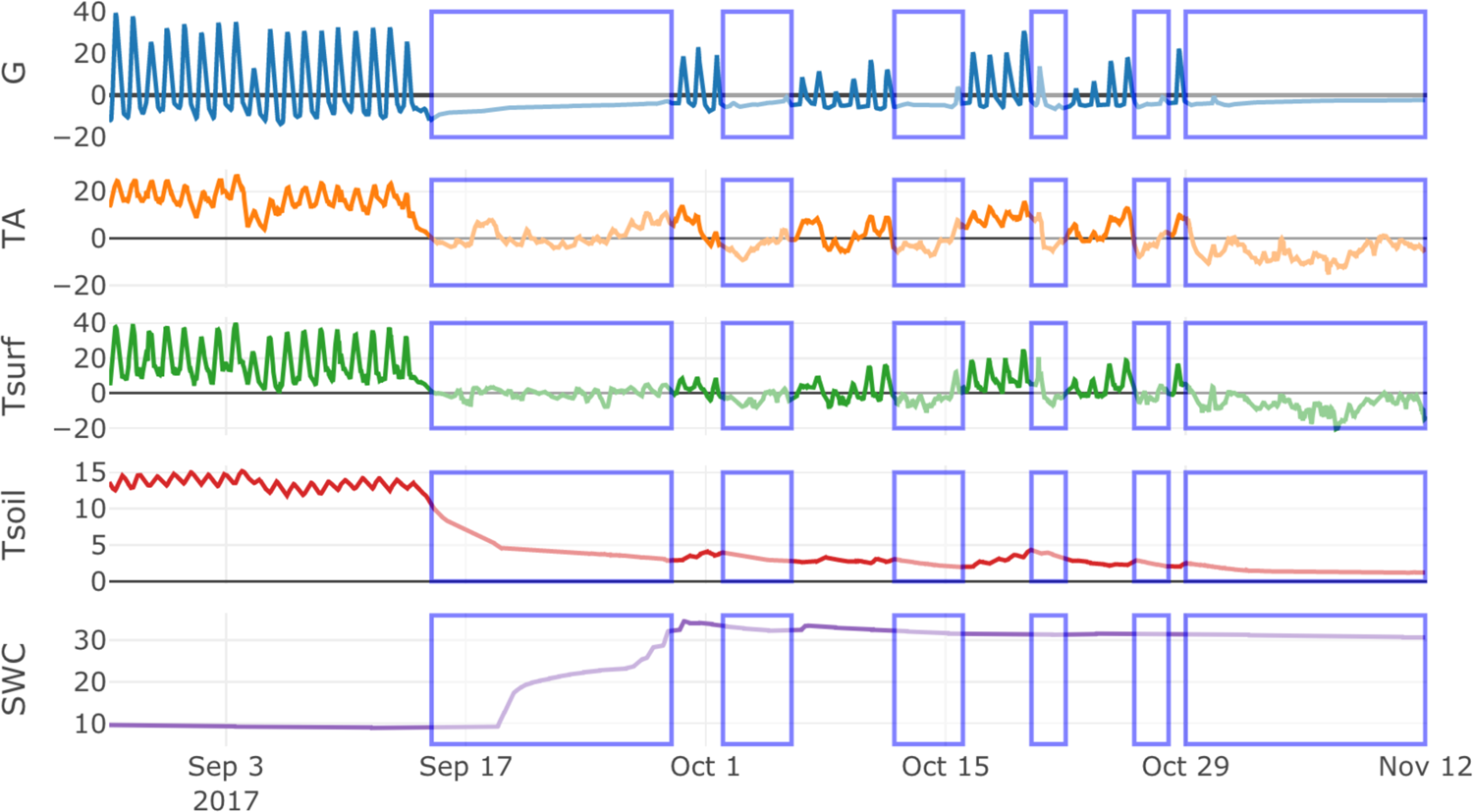
The same as Figure 2 but for 2017. Periods identified as having snow (Table 3) are indicated in blue boxes.

**Figure 7:**
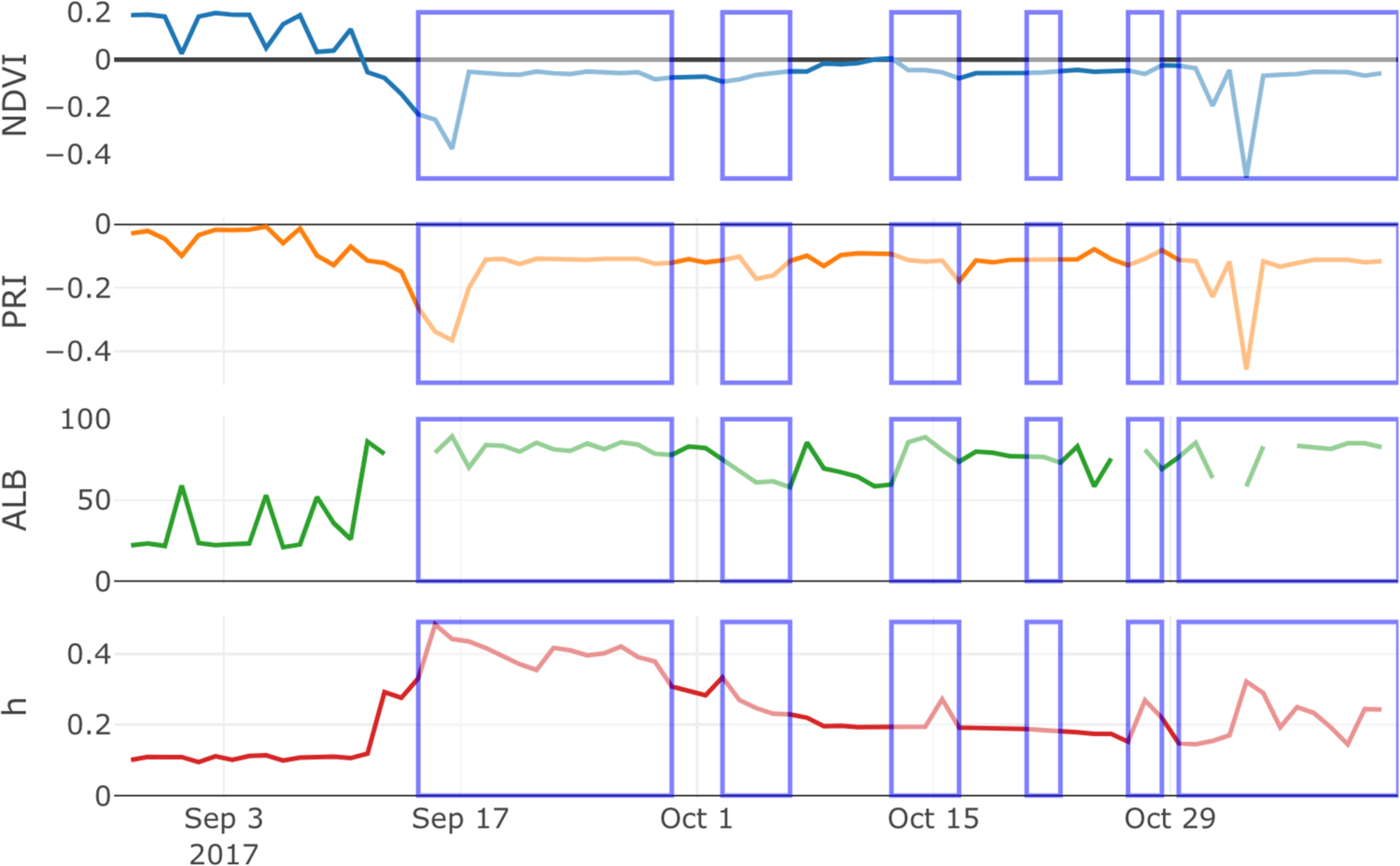
The same as Figure 3 but for 2017. Periods identified as having snow (Table 3) are indicated in blue boxes.

**Figure 8:**
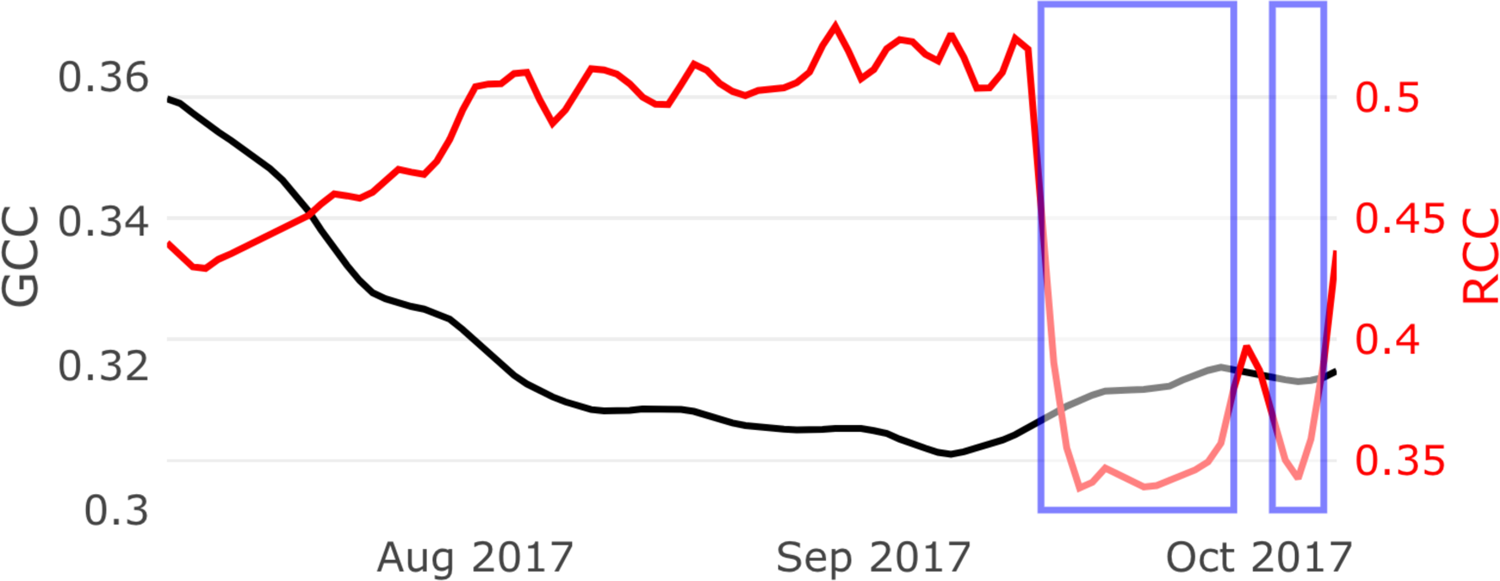
The smoothed daily 90th percentile green chromatic coordinate (GCC) and red chromatic coordinate (RCC) from phenocam observations at the Bangtail Mountain (MT, USA) study site during 2017 for periods when the phenocam was active. Periods of snow identified using visual analyses of phenocam images are indicated in blue boxes (Table 3).

**Table 3:**
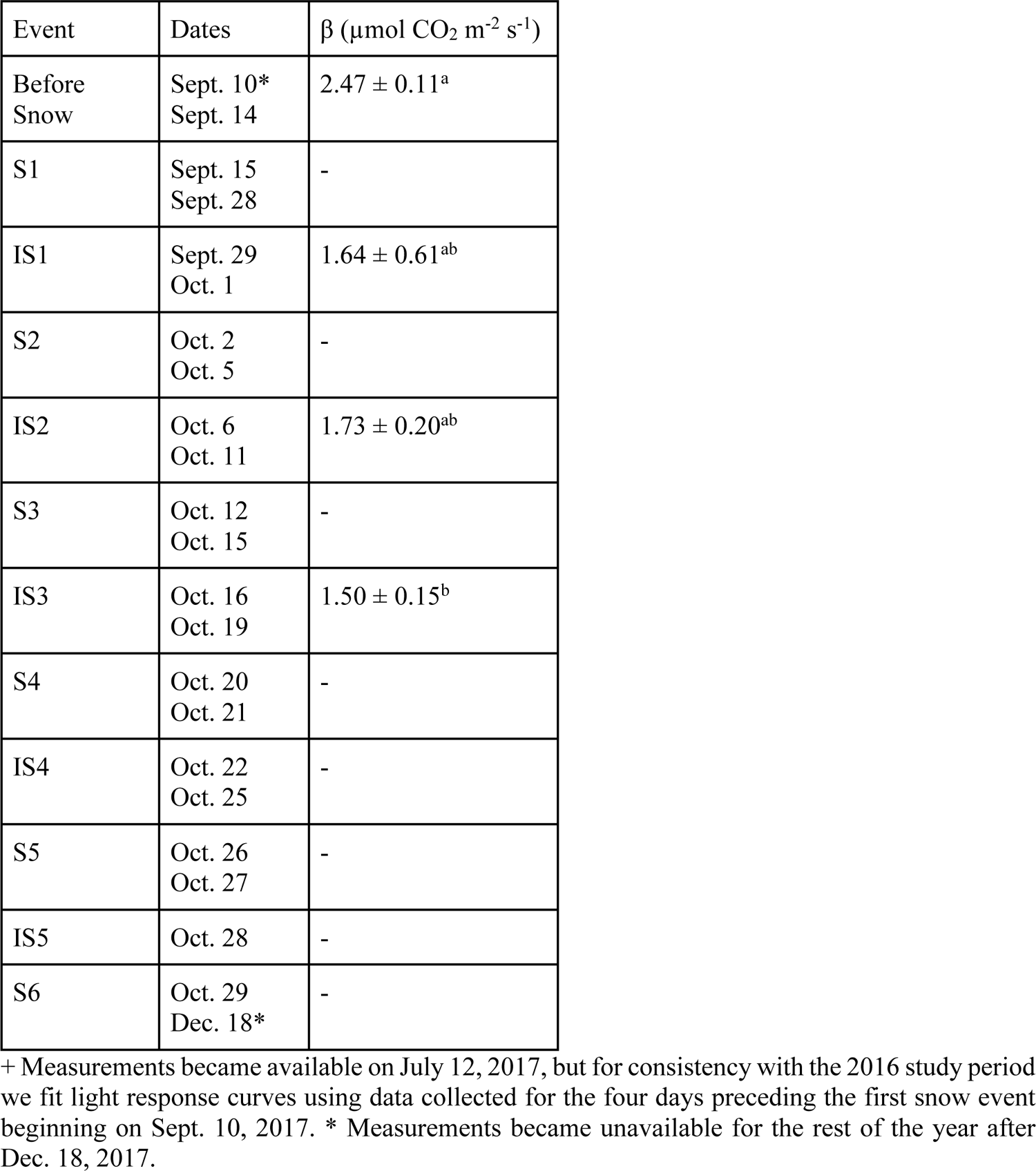
The same as Table 2 but for 2017. A light response curve was not calculated for the single day of available data during IS5, and the data were not significantly related to the light response curve during IS4.

| Event | Dates | $\beta$ ($\mu\text{mol CO}_2 \text{ m}^{-2} \text{ s}^{-1}$) |
| --- | --- | --- |
| Before Snow | Sept. 10*<br>Sept. 14 | $2.47 \pm 0.11^a$ |
| S1 | Sept. 15<br>Sept. 28 | - |
| IS1 | Sept. 29<br>Oct. 1 | $1.64 \pm 0.61^{ab}$ |
| S2 | Oct. 2<br>Oct. 5 | - |
| IS2 | Oct. 6<br>Oct. 11 | $1.73 \pm 0.20^{ab}$ |
| S3 | Oct. 12<br>Oct. 15 | - |
| IS3 | Oct. 16<br>Oct. 19 | $1.50 \pm 0.15^b$ |
| S4 | Oct. 20<br>Oct. 21 | - |
| IS4 | Oct. 22<br>Oct. 25 | - |
| S5 | Oct. 26<br>Oct. 27 | - |
| IS5 | Oct. 28 | - |
| S6 | Oct. 29<br>Dec. 18* | - |

+ Measurements became available on July 12, 2017, but for consistency with the 2016 study period we fit light response curves using data collected for the four days preceding the first snow event beginning on Sept. 10, 2017. \* Measurements became unavailable for the rest of the year after Dec. 18, 2017.

Maximum daily GPP before and after the first two snow events was on the order of 3 μmol m^-2^ s^-1^ (Figure 9); recall that light-saturated GPP (i.e., β) reached values of ∼6 μmol m^-2^ s^-1^ in 2016 (Figures 4 and 5). β was 2.47 μmol m^-2^ s^-1^ before snowfall. Like 2016, β was not significantly different for the first two inter-snow periods but was significantly less than the before-snow period by the third inter-snow period, when it decreased to 1.50 μmol m^-2^ s^-1^ (Figure 10, Table 2). As opposed to 2016, NEE was rarely less than 0 μmol m^-2^ s^-1^ such that the ecosystem was almost continuously losing carbon after the first snowfall in 2017 (Figures 4 and 9). The ecosystem was a consistent net source of C to the atmosphere as a consequence, with the possible exception of two relatively large net carbon uptake events in early November that are inconsistent with micrometeorological observations during this time that suggest snow presence. The ecosystem lost 51 g C m^-2^ during the period from October 1 - November 12, 2017, similar to 2016. There was a notable increase in NEE and RE from values less than 2 μmol m^-2^ s^-1^ to values greater than 2 μmol m^-2^ s^-1^ around September 17, 2017, a snow-covered period when TA was greater than 0 ℃, SWC increased from 10 to 20%, LE reached values of nearly 100 W m^-2^, and H was negative during the day indicating heat flux into the snowpack (Figures 6 and 9).

**Figure 9:**
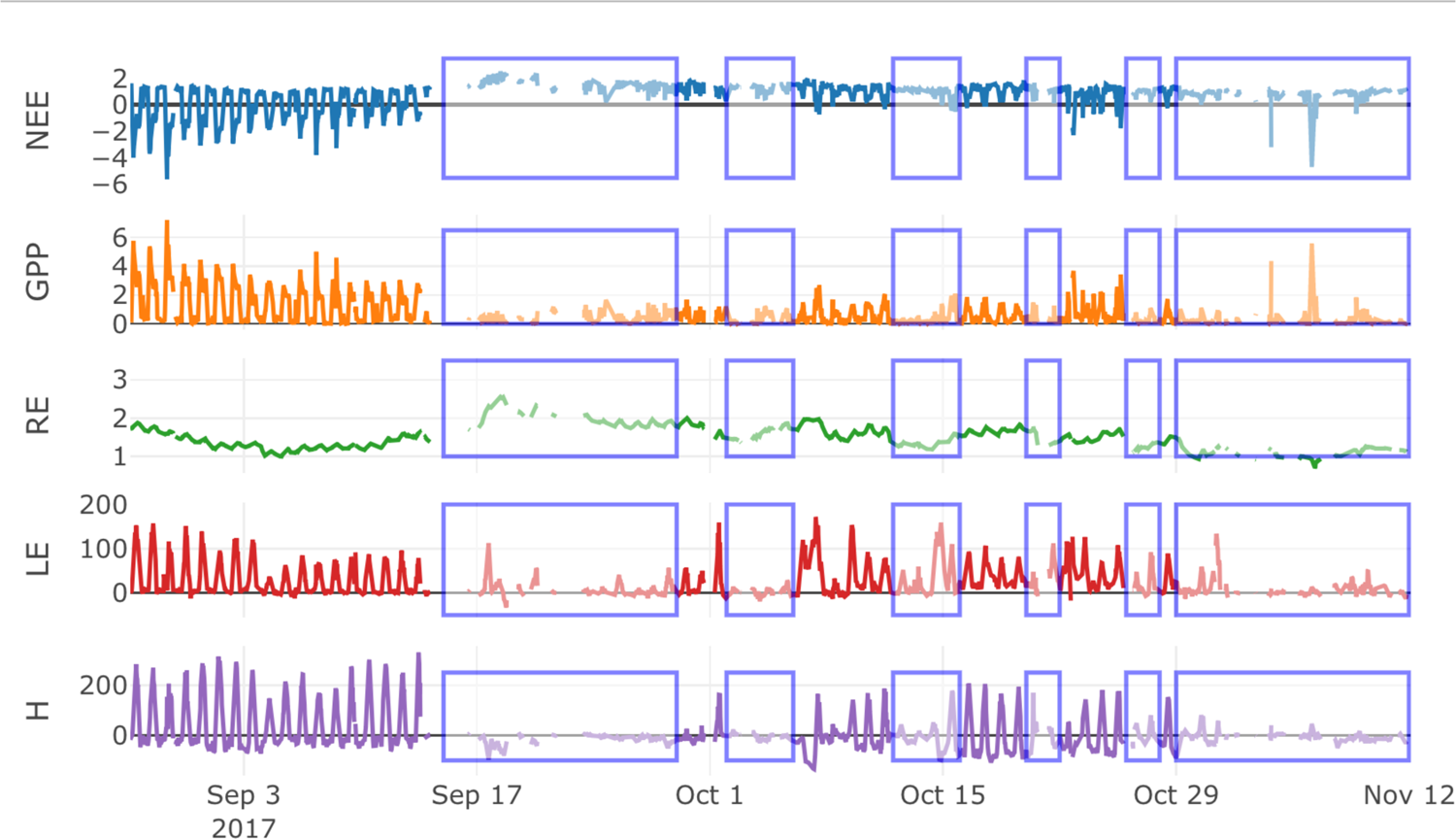
The same as Figure 4 but for late August through mid-November 2017. Periods identified as having snow (Table 3) are indicated in blue boxes.

**Figure 10:**
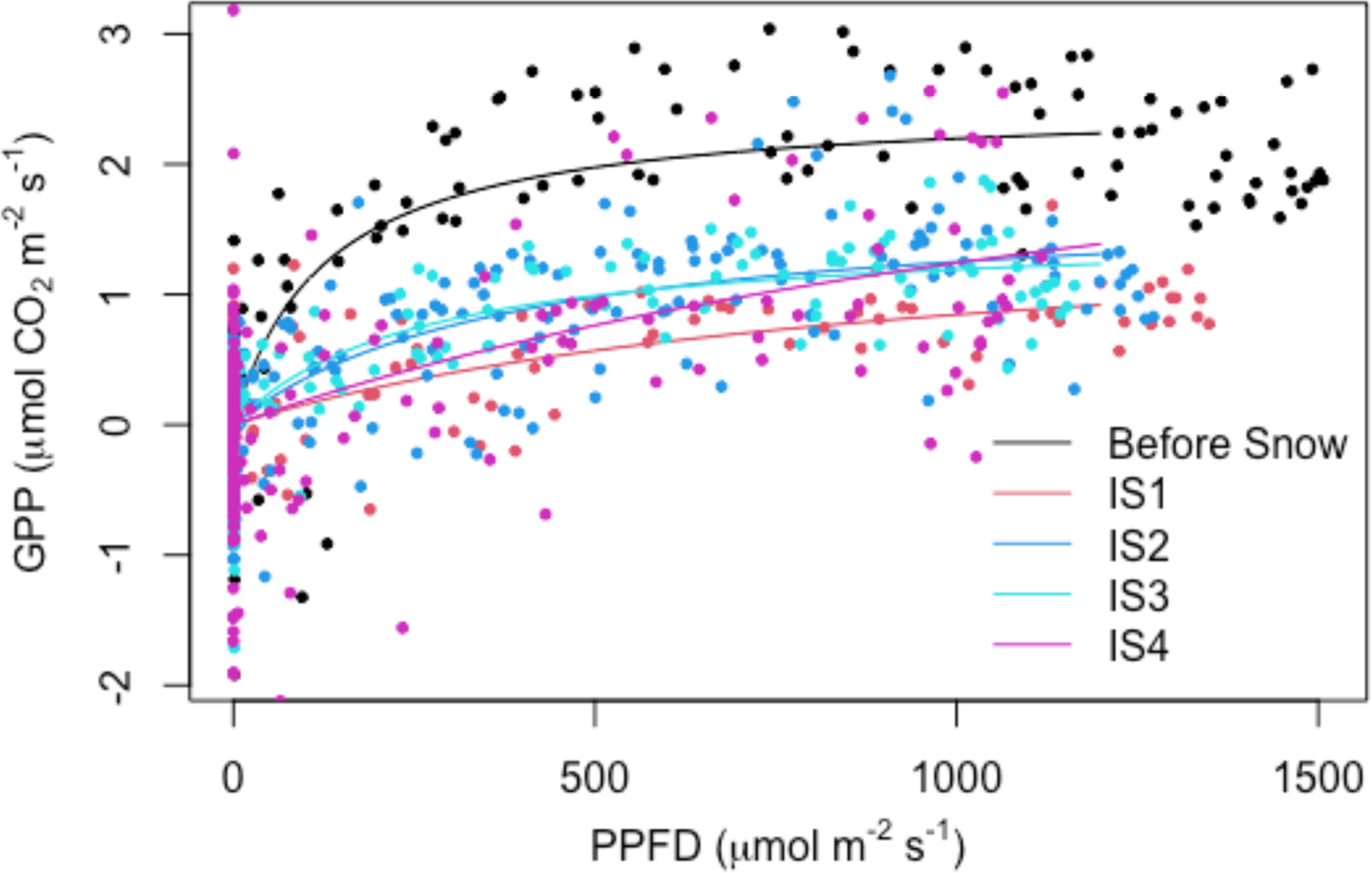
The same as Figure 5 but for 2017 with inter-snow periods (IS) as described in Table 2.

### Observations during summer, 2017

Eddy covariance and micrometeorological measurements were available from mid-July 2017 onward (Figure 11) and we present these observations to place the 2017 autumn measurements in context. Light-saturated GPP (i.e., β) was 18.6 μmol m^-2^ s^-1^ during late July (Figure 12), already decreased to 9.02 μmol m^-2^ s^-1^ as drought progressed in early August, decreased further to 6.16 μmol m^-2^ s^-1^ during late August, and reached ∼3 μmol m^-2^ s^-1^ during early September as noted above (Figure 10, Table 2). In other words, β was approximately 1/6^th^ of its maximum observed value in summer before snowfall in 2017.

**Figure 11:**
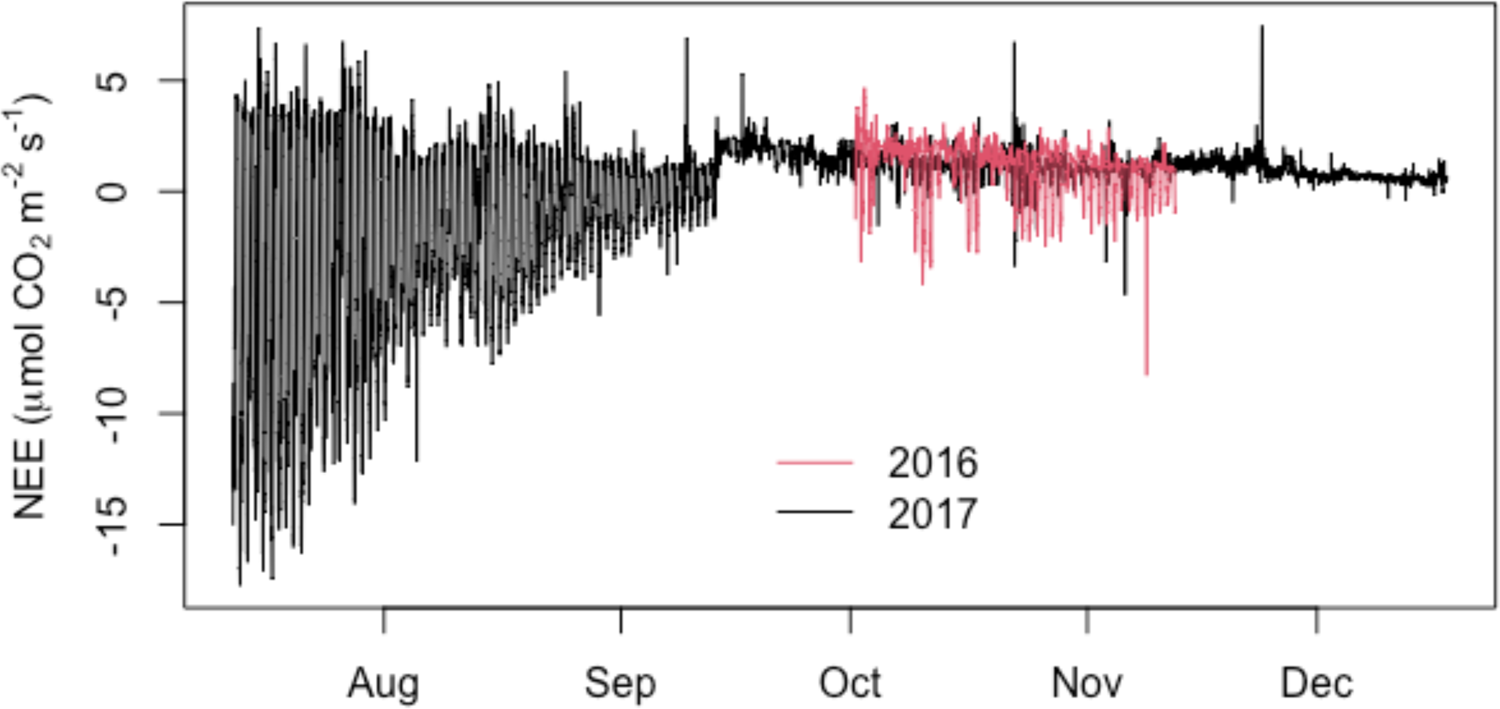
The net ecosystem exchange of carbon dioxide (NEE) measured during 2016 and 2017 at the Bangtail Mountain Meadow research site. Time series are constrained by measurement availability and values represent measurements and gapfilled estimates using REddyProc (Wutzler et al., 2018).

**Figure 12:**
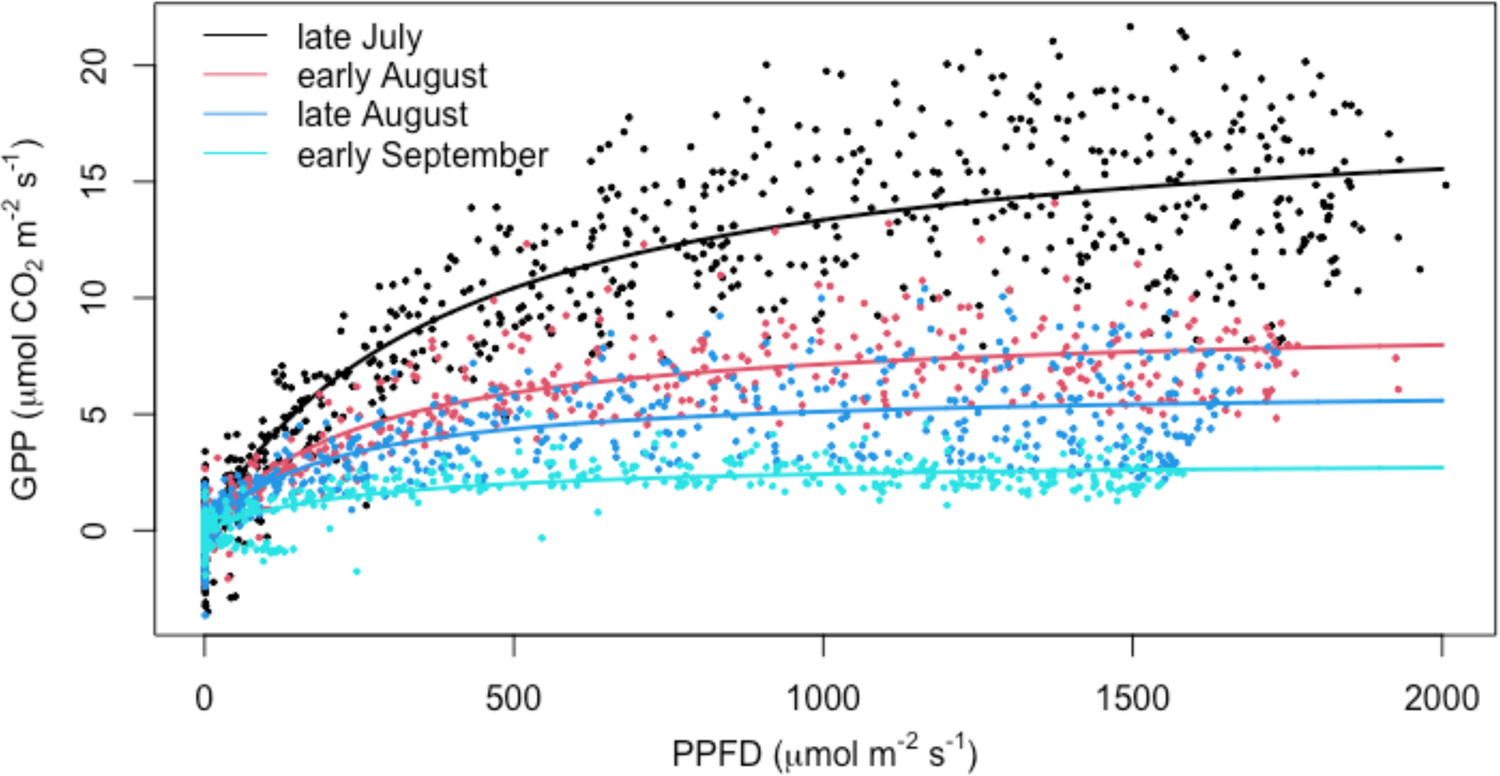
The relationship between GPP and PPFD for the periods July 12-31 (‘late July’), August 1-14 (‘early August’), August 15-31 (‘late August’), and September 1-15 (‘early September’), 2017 with fitted light response curves following Figure 5.

#### MODIS

We analyzed only those NDVI_MODIS_ observations greater than 0.4 to help avoid noisy measurements at lower values that are consistent with full or patchy snow cover. A notable increase (i.e., greening) in NDVI_MODIS_ occurred in 2015 from 0.45 on October 7 to 0.5 on October 27, and NDVI_MODIS_ approached 0.5 in 2017 after a long period of low NDVI_MODIS_ values with an inconsistent vegetation signal that began on September 12 (Figure 13). Other years, namely 2016 and 2019, did not demonstrate any late season increase in NDVI_MODIS_ although measurements in 2016 rarely dropped below 0.4 during the latter part of the calendar year.

**Figure 13:**
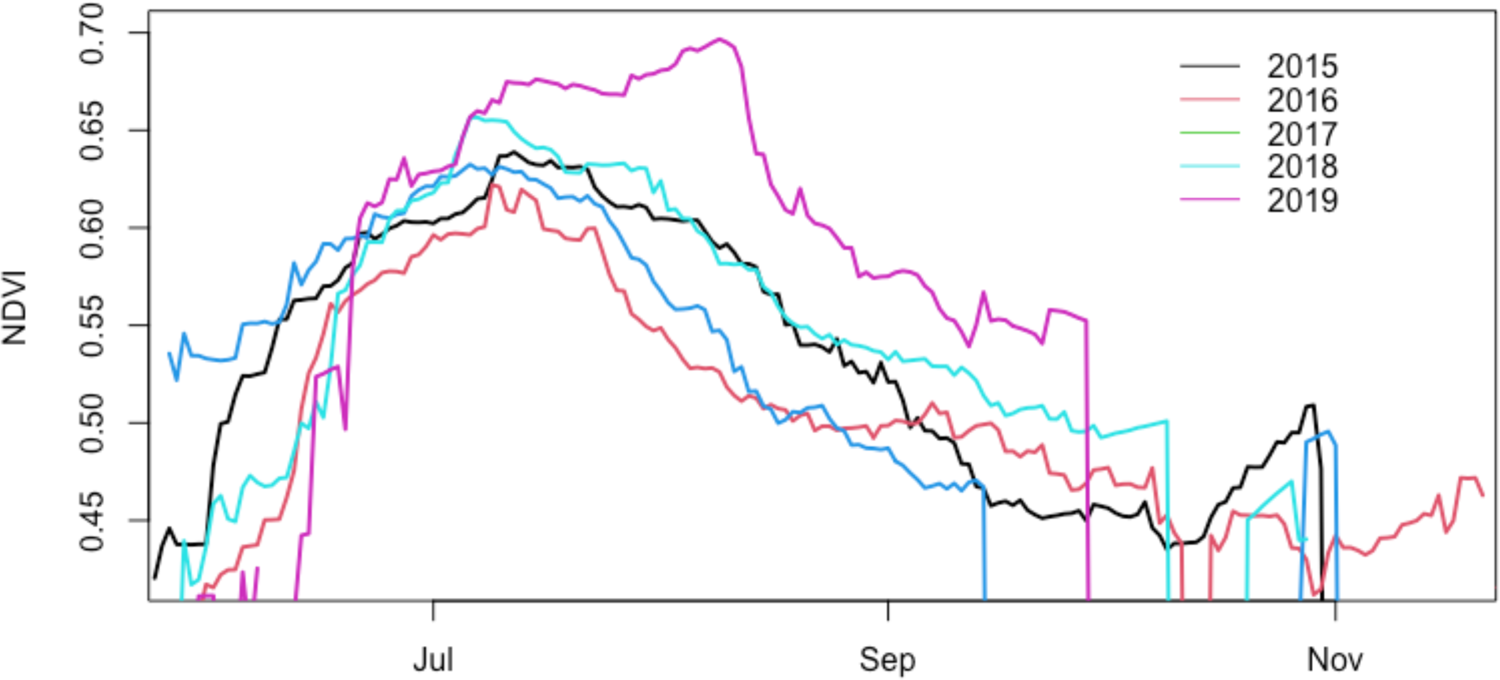
MODIS NDVI for the Bangtail Mountain Meadow study site from the MCD43A4 v006 500 m daily product.

## Discussion

*Overview: Is photosynthesis in mountain meadows resistant or resilient to autumn snow events?* Decades of research at the Bangtail Mountain Meadow study site and communication with the Montana ranching community has led to the suggestion that canopy photosynthesis would “recover” after drought-breaking autumn precipitation, which often falls as snow. The ecosystem was not likely under drought stress before snow fell in 2016 as SWC was ∼ 30% (Figure 3) but was only ∼10% before snowfall during 2017 (Figure 7) and thus well below the point at which grasses are often considered drought stressed (Rodríguez-Iturbe and Porporato 2007; Rodriguez-Iturbe et al. 1999). Observations from 2017, a drier year, indicate that GPP was rarely greater during inter-snow periods than it was before snowfall despite added moisture (Figures 4 and 9), and β was not significantly different before or after the first two snow events in either year (Figures 5 & 10, Table 2). Observations from both years also do not necessarily suggest that there is a lag in the recovery of GPP after snow melt (Figures 4 and 9). In other words, rather than a recovery of photosynthetic function and thereby *resilience* to snow disturbances, our observations suggest that the vegetation probably never lost photosynthetic function under snow in the first place, at least after the first two snow events. Instead, observations are consistent with the notion that autumn snowfall and melt is a minor disturbance against which mountain meadow photosynthesis is *resistant*.

Temperate, boreal, and arctic grasses can photosynthesize at temperatures at or below 0 °C (Tieszen 1973; Tieszen and Sigurdson 1973; Skinner 2007), and some grasses maintain photosynthesis under cold and dark conditions that simulate the subnivean environment (Höglind, Hanslin, and Mortensen 2011). Our observations suggest finite GPP even in the presence of snow (e.g., Figures 4 and 9). At the same time the study site is vegetatively diverse, and it is difficult to establish with certainty that grasses are the primary contributor to ecosystem-scale GPP during and after snow events. Some mosses can photosynthesize under snow (Street et al. 2012) and are characterized by light-saturated photosynthesis (e.g., β) ranging from nearly 0 to 7 μmol m^-2^ s^-1^ (Van Gaalen, Flanagan, and Peddle 2007; Street et al. 2012; Skre and Oechel 1981), similar to our observations during periods between snow events in 2016 (Figure 5). At the same time, moss cover at the site is sparse and likely insufficient to explain β observations of up to 7 μmol m^-2^ s^-1^ on a ground area basis. Instead, photosynthesizing vascular vegetation – that often occurs toward the bottom of grass canopies in autumn (Migliavacca et al. 2011) – is more consistent with available observations (see Figure A1). These patterns are often difficult to discern with phenocams that point laterally (Figures 1 and 8) (Migliavacca et al. 2011). The tower-mounted NDVI sensors point downward, and measurements can exceed 0.2 during periods between snow (e.g. Figure 2). These are low values of NDVI for grass sites (Gamon et al. 1995) that are consistent with the small carbon fluxes that we observe during these times (Figures 4, 5, 9, and 10). Carbon uptake is also lower at the study site than most global grasslands; for example, peak light-saturated NEE at 1500 μmol m^-2^ s^-1^ PPFD following the approach of (Ruimy et al. 1995) is 10.9 μmol m^-2^ s^-1^ at our study site, versus a global average of 25.5 μmol m^-2^ s^-1^ (Gilmanov et al. 2010), as would be expected for our relatively cold study site.

Our measurements gave no evidence that mountain meadow GPP was any greater after snow than before snow, even when more moisture and favorable growing conditions occurred after snowmelt periods as seen in 2017. It was unclear if there was any flush of new leaves, but no observable height growth between snow events occurred (Figures 3 and 7). We cannot say for certain which grass species were responsible for photosynthesis after snow, but observations in Bozeman suggest that the invasive *Poa pratensis* rather than native *Festuca idahoensis* is more likely to remain green in the cold season, with consequences for ecosystem function as invasive grasses continue to establish at the study site. Species-level measurements at the Bangtail Mountain Meadow would likely require careful in-person measurements during uncertain travel conditions and/or high-resolution imagery that is increasingly used in conjunction with machine learning approaches to estimate grass growth and phenology (Oliveira et al. 2020; Viljanen et al. 2018). A comprehensive monitoring framework would require species-level studies of the response of the ecosystem to climate variability and would help reconcile population and community dynamics that we were unable to observe with certainty in the present study.

### Ecosystem dynamics during snow presence

A notable event occurred around September 17, 2017, when snow was present, but TA reached nearly 10 ℃ and H became negative, indicating heat exchange into the snowpack and melt (Stoy et al. 2018), but Tsurf remained at or near 0 ℃ indicating that snow did not fully melt. SWC increased from 10% to 20% during this event and NEE also increased, suggesting enhanced soil respiratory activity. We posit that the snowpack was melting during this period but did not fully ablate, and the corresponding input of soil water enhanced respiratory activity owing to the Birch effect (Jarvis et al. 2007). We further speculate that the soil water input and observations of respiratory fluxes were synchronized in time because of pressure pumping (i.e., enhanced diffusion) through snow at this characteristically windy montane site (Rains et al. 2016; Bowling and Massman 2011; Berryman et al. 2018). Decadal snow fence observations at the site that suggest that enhanced snowpack duration depletes soil C, N and P (Yano et al. 2015; Weaver and Collins 1977), but the microbial mechanisms of potential under-snow Birch effects in ecosystems that receive drought-breaking snow need to be substantiated with direct observations of soil biogeochemistry.

### The challenges of patchy snow

We used micrometeorological and radiometric observations to identify snow presence using instruments on or in the vicinity of the eddy covariance tripod tower, confirmed with phenocam measurements when available (Figures 1, 2, 3, 6, 7 and 8). Turbulent fluxes arise from the eddy covariance flux footprint, which was often in the scene of the phenocam that measured the area immediately to the west of the tower, the dominant wind direction (Figure 1). The physical infrastructure of the tower will cause snow deposition and alter snowpack characteristics; and we suspect that this is the reason that the 2017 melt events identified using micrometeorological and flux observations (Figures 6 and 9) differed from the radiometers that suggested snow was still present (Figure 7). In other words, snow presence was often patchy (e.g., Figure 1) and influenced by the measurement infrastructure itself, by slowing wind speed to deposit snow. This leads to an observational challenge of identifying what is ‘snow-covered’ for the purpose of ecosystem-scale studies. Going forward, we propose a tripartite classification system with no snow, full snow, and patchy snow may be the best approach for studying the impacts of snow at the plot or flux footprint scale, which remains a challenge for observation systems (Kosmala et al. 2018). During melt, dynamic changes to the snow-covered area across space and time are being sampled by a dynamic flux footprint (Chu et al. 2021), both of which require extensive quantification that can include substantial uncertainty. From our analysis such an observation system could not rely on tower-mounted instrumentation alone due to tower infrastructure impacts on snow dynamics. In this case a tower within the field of view of the phenocam would have been preferred (Stoy et al. 2021).

### Remote sensing

Remote sensing plays an important role in understanding the physiology of global ecosystems, especially in areas that are difficult to access. MODIS has been found to provide reasonable measurements of vegetation physiology for grasslands of the Northern Rockies (St. Peter et al. 2018). Differences are to be expected amongst RS sensors (Rossi et al. 2019) and among different grassland sites (Wohlfahrt et al. 2010), and the coarse spatial scale and diurnal return interval of MODIS gave very different NDVI results than the tower-mounted instrumentation at our site (Figures 3, 7, and 13). Recent advances to geostationary satellite technology – for example enhanced observational capacities in the visible and near infrared from the Advanced Baseline Imager on the Geostationary Operational Environmental Satellite (Khan et al. 2021) – provide new opportunities for understanding snow duration (Romanov and Tarpley 2004; Romanov et al. 2000) and vegetation production after melt. Enhanced observational systems at multiple scales in space including tower-mounted instruments, phenocams, unpiloted aerial vehicles, and satellite remote sensing will further improve our understanding of autumn phenological changes in global ecosystems (Gallinat et al. 2015).

### Implications and future studies

Grass photosynthesis is notably resilient to small disturbances. If sufficient soil moisture is available for regrowth photosynthesis tends to reestablish rapidly following harvesting (Novick et al. 2004; Stoy et al. 2008), grazing (Parsons et al. 1983), and heat waves (Hoover, Knapp, and Smith 2014), often dependent on species and growth form (Cremonese et al. 2017). Grass photosynthesis also tends to establish rapidly after snowmelt in spring (Julitta et al. 2014; Vitasse et al. 2017), which we were not able to observe at our site due to equipment and power failure over winter. However, our observations suggest that mountain meadow photosynthesis is resistant to autumn snowfall which appears to be a minor disturbance for events up to two weeks in duration. The resistance of grass vegetation to extreme climate conditions is another matter as extreme drought (Hoover, Knapp, and Smith 2014) and desertification (D’Odorico et al. 2013) threaten grasslands worldwide.

It has long been known that the snowpack dynamics are a key determinant of the phenology and productivity of subalpine meadows (Canaday and Fonda 1974). We argue that spring snowmelt (or summer, depending on site) is only one of often two periods of importance for montane grassland phenology and function, the second being the transition to the full snowpack in winter, which may contain multiple snowfall and melt periods that can bring needed moisture to mountain meadows (Brookshire and Weaver 2015). Montane grass photosynthesis appears to be resistant at the ecosystem-scale, at least for the first two snow events in our study. Enhanced observational systems that consider population, community, and ecosystem-scale responses to ongoing climate changes during autumn (Gallinat et al. 2015) will improve our understanding of the ecosystem services that montane grasslands provide across all seasons.

## Acknowledgments

We acknowledge the Indigenous Nations on whose ancestral lands the study took place and recognize that the infrastructure used for this project was built on Indigenous land. We recognize multiple Indigenous Nations as past, present, and future caretakers of this land whose stewardship of the region was interrupted by their physical removal under the 1830 Indian Removal Act and through US Assimilation policies explicitly designed to eradicate Indigenous language and ways of being until the 1970s. This work was supported by the National Science Foundation under the EPSCoR Track II cooperative agreement OIA-1632810. We also thank the Montana State University College of Agriculture and the University of Wisconsin for financial assistance and Dr. James Irvine for extensive technical support. Adam Cook, Dr. Geneva Chong, Justin Gay, Dr. Bill Kleindl, F. Aaron Rains, Dr. Angela Tang, and Bill Vandenberg provided additional technical support for which we are grateful., And we thank Dr. Susanne Wiesner for insightful comments on the manuscript. Phenocam research was supported by Colorado State University and the AmericaView program (grants G13AC00393, G11AC20461, G15AC00056) with equipment and deployment sponsored by the Department of Interior North Central Climate Science Center.

---

**Figure A1:**
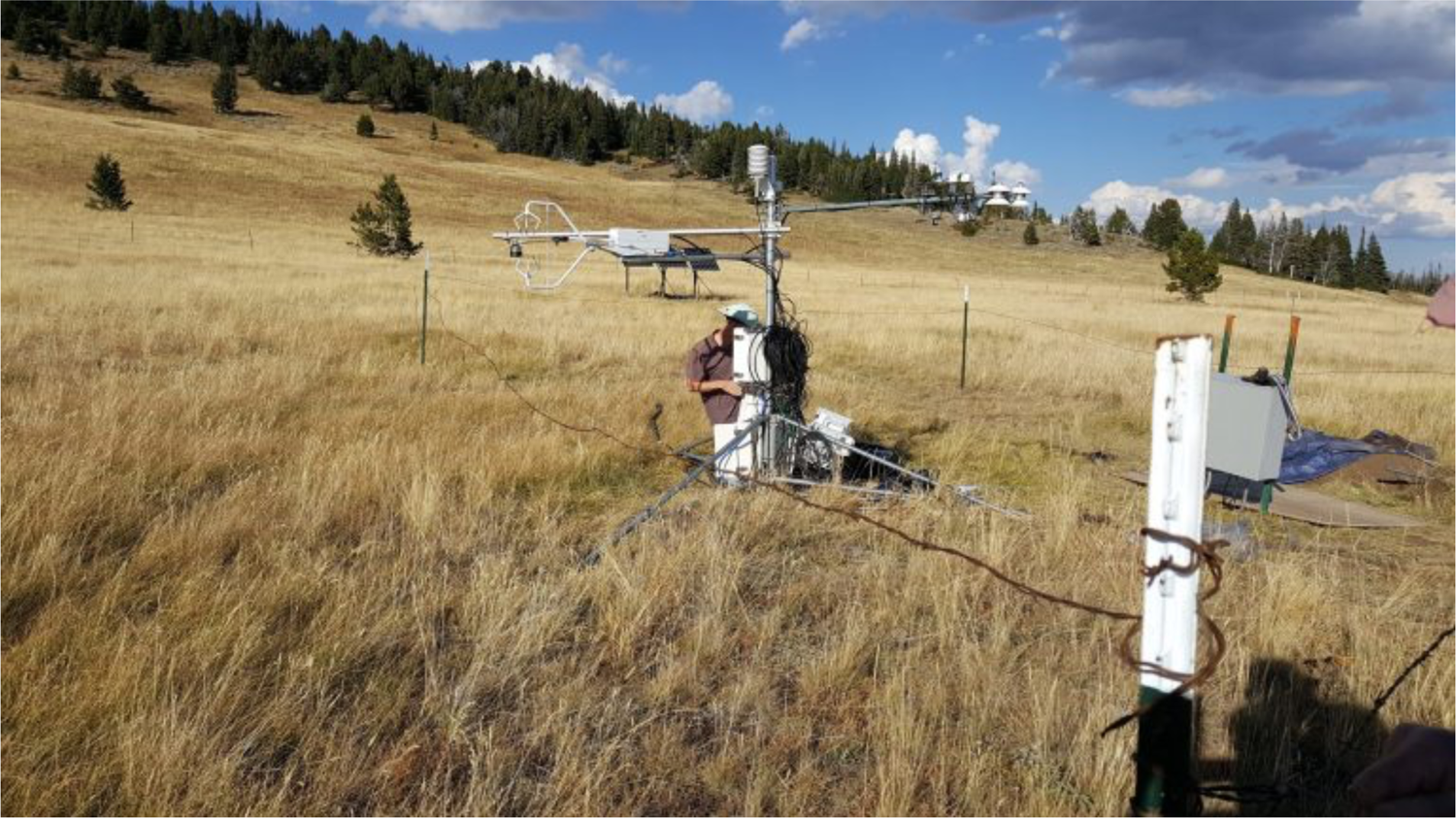
The Bangtail Mountain Meadow eddy covariance tower (US-BMM) during installation on September 30, 2016.

**Figure A2:**
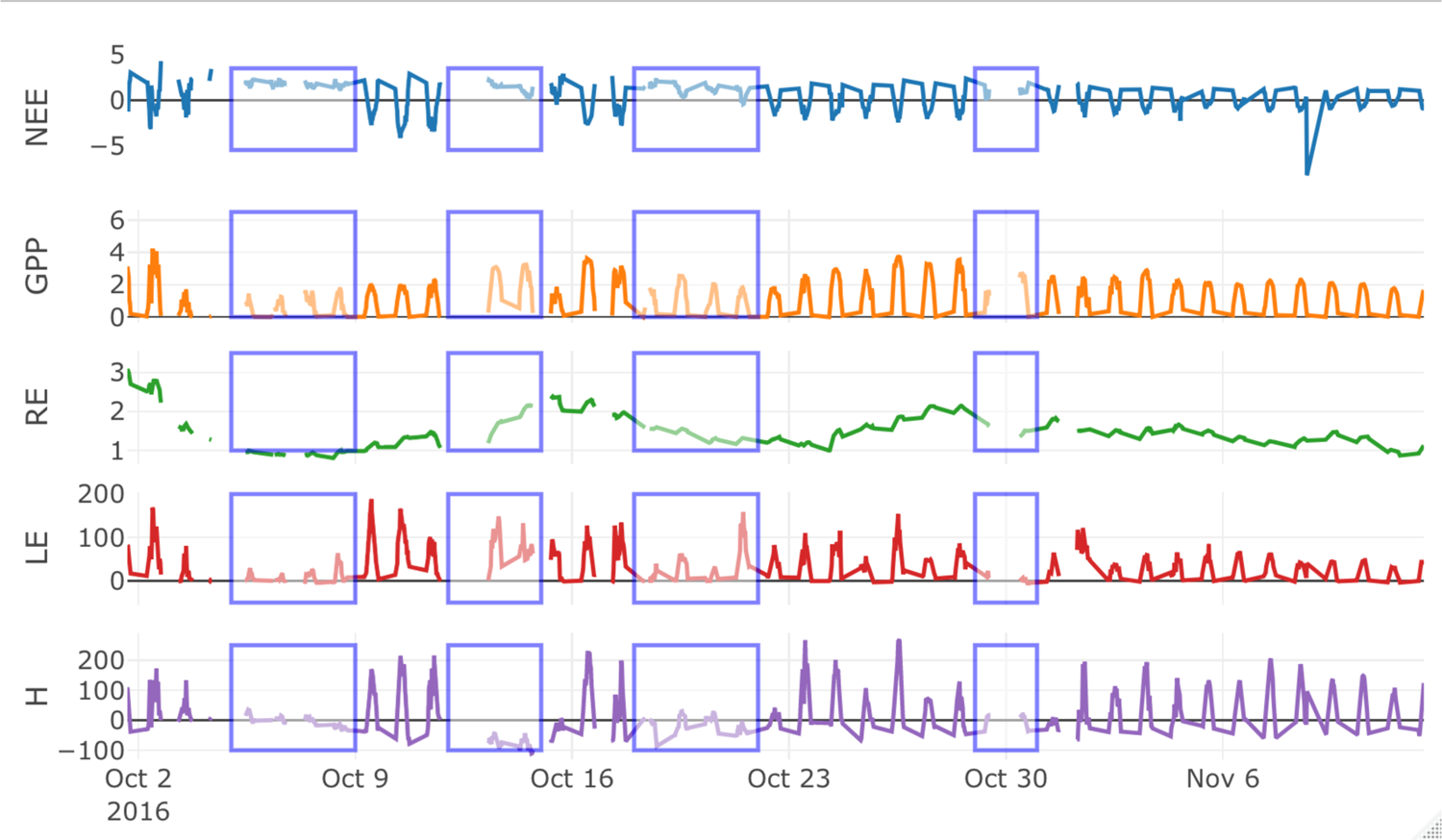
The same as Figure 4 but using the Lasslop et al. (2017) daytime flux partitioning approach.

